# Targeted expansion of Protected Areas to maximise the persistence of terrestrial mammals

**DOI:** 10.1101/608992

**Authors:** Sarah Mogg, Constance Fastre, Martin Jung, Piero Visconti

**Author notes:** **Corresponding authors:** Piero Visconti.

## Abstract

Over a quarter of species assessed by the IUCN Red List are threatened with extinction. A global commitment to protect 17% of land and 10% of the oceans by 2020 is close to being achieved, but with limited ecological impacts due to its inadequacy and poor enforcement. Here, we reverse-engineer IUCN Red List criteria to generate area-based conservation targets and spatial conservation priorities to minimize the extinction risk of the world terrestrial mammals. We find that approximately 60% of the Earth’s non-Antarctic land surface would require some form of protection. Our results suggest that global conservation priority schemes, among which the Aichi targets, will be inadequate to secure the persistence of terrestrial mammals. To achieve this goal, international cooperation is required to implement a connected and comprehensive conservation area network, guided by high priority regions outlined in this study.

## INTRODUCTION

In recent decades global biodiversity has undergone increasing threat from anthropogenic activities (Tittensor et al. 2014; Joppa et al. 2016; Maxwell et al. 2016). Today, approximately 27% of assessed species are at risk of extinction (IUCN 2018) and this figure is predicted to increase (Newbold et al. 2015; Visconti et al. 2016). Recognizing the urgency for rapid action, the parties of the Convention on Biological Diversity (CBD) committed to the Strategic Plan for Biodiversity 2011-2020 and its 20 Aichi Biodiversity Targets (ABT). Rooted on the potential of effectively managed protected areas (PAs) as critical conservation tool (Geldmann et al. 2013; Watson et al. 2014), Aichi Target 11 advocates for the conservation of 17% of terrestrial and 10% of marine environments worldwide, particularly areas of global significance to biodiversity, into a connected network of such PAs or other area-based conservation measures (CBD 2010). This target should contribute to achieve Aichi target 12 which aims to prevent the extinction and improve the conservation status of known threatened species (CBD 2010).

Recent studies have criticized Aichi Target 11 for its ecological inadequacy and lack of ambition (Venter et al. 2014), for its ambiguity (Butchart et al. 2016) and its vulnerability to the “gamed” thereby producing perverse outcome (Barnes et al. 2018). In particular, promoting the conservation of large protected areas of little conservation value to achieve the coverage percentage element of the target, comes at the expense of producing biodiversity impacts (Pressey et al. 2015). As a result, the more ambitious proposal of setting aside Half-Earth for biodiversity has gathered support among conservationists (Noss et al. 2012; Wilson 2016; Watson & Venter 2017) but its potential to deliver positive biodiversity outcomes remains untested. Therefore, the question remains as to how much land is required, and where these should be placed, to achieve species conservation.

So far, the proportion of a species’ range to be targeted for protection has been set arbitrarily (Rodrigues et al. 2004; Ceballos et al. 2005; Carwardine et al. 2008). Even though the method introduced by Rodrigues et al. (2004) for mammals has been adopted in other studies (Venter et al. 2014; Butchart et al. 2015; Visconti et al. 2016), it may be inadequate to reduce the extinction risk of the species. We therefore propose a conservation target setting approach based on the thresholds used in the IUCN Red List criteria (Box S1, IUCN 2012). We use these targets to identify global area-based conservation targets required to maximize species persistence (Aichi Target 12). By deriving area-based conservation targets based on population ecology and extinction risk analyses, this study aims to address the following questions: (1) For terrestrial mammal species, determine the geographic extent requiring protection to maximize their long-term persistence, as informed by the IUCN Red List criteria; (2) On a global scale, identify critical regions in which area-based conservation strategies could be expanded to encompass the above targets.

## METHODS

### 1. Study species

This study focused on global terrestrial mammals assessed and classified under the IUCN Red List with ranges downloadable from www.iucnredlist.org. We focus on mammals to keep the analysis computationally tractable and their distribution is generally well described and diversity patterns in mammals are overall representative of other major taxonomic groups (Qian and Ricklefs 2008). After excluding terrestrial water-dependent species and those with no range data or information on habitat preferences, a remaining 4325 terrestrial mammal species were considered.

### 2. Conservation objectives

#### 2.1 Targets informed by extinction risk criteria

We aimed to set IUCN-informed target areas that would ensure that each species has an Area Of Occupancy (AOO, the area which is occupied by a taxon, excluding cases of vagrancy, IUCN 2012), sufficiently large to qualify for IUCN Red List category “Least Concern”. To achieve that, based on the RL criteria, two main thresholds were considered for target-setting: (1) Based on criterion A, a species’ population must not decline more than 30% within 10 years or three generations (whichever the longer), to avoid being classified as “Vulnerable”. Allowing for a 10% buffer (generally applied to separate the Least Concern from the Near Threatened category), and assuming a linear relationship between changes in population and changes in species’ range, the species’ target area must therefore not be below 80% of a species’ range. (2) Based on criterion B2, a species’ AOO must not fall below 2,000 km^2^. With a 10% buffer, a species’ target area must therefore not fall below 2,200 km^2^. Following Butchart, et al. (2015) we applied an upper limit of 1,000,000 km^2^ for all species with ranges greater than 1,250,000 km^2^, due to the logistic difficulties in creating extremely large PAs. Target areas were therefore determined as 80% of a species’ range, with 2,200km^2^ and 1,000,000 km^2^ as the lower and upper limits, respectively.

We calculated two variants of these IUCN-informed targets: one variant based on the species range size (RSI targets) and a variant based on the suitable habitat available within the species range (HSI targets). RSI targets were produced by applying the criteria above to the native and extant portion of terrestrial mammal range maps available from the IUCN Red List database, accessed in June 2018 (IUCN 2018) for comparison with previous targets that were designed to work with and were applied to this data (*e.g.* Rodrigues et al. 2004; Venter et al. 2014; Butchart et al. 2015). However, species generally do not occupy the full extent of their range and applying persistence targets based on AOO to range maps may falsely assume that conserving any part of a species range would contribute to its persistence. The Extent of Suitable Habitat (ESH), which results from subtracting all habitat types considered unsuitable (according to the IUCN Red List species accounts) from the species’ range may thus constitute a better proxy for the AOO of each species as it reduces the commission error relative to using range maps (Rondinini et al. 2011). We calculated the HSI targets using the ESHs produced for each species using IUCN ranges as a base map and land-cover and land-use data reconstructed for the year 2015 from the IMAGE modelling platform (Stehfest et al. 2014), details of the data and methods are described in Visconti et al. (2016).

#### 2.2 Targets informed by range size

To compare our IUCN-based range-size targets (RSI targets) with the targets applied in several global conservation planning studies, we reproduced the range-size targets initially proposed by Rodrigues et al., 2004 (RS targets), using the expert-based geographic ranges from the IUCN Red List database mentioned above. We assigned targets equating 10% and 100% of their range to widespread (range > 250,000 km^2^) and small-ranging (range < 1,000 km^2^) species, respectively. For other species, we applied log-linear interpolation between these percentages.

### 3. Gap in species coverage by current protected areas

We assessed the extent to which species’ ranges were sufficiently covered by protected areas based on each of the 3 targets, their range and the current protected area estate (WDPA updated to April 2019). We did this for the world terrestrial mammals, birds and amphibians.

### 4. Determining potential conservation area networks

We refer to conservation area networks as any area that could be targeted for habitat retention and biodiversity conservation, and therefore contribute to the goals of Aichi Targets 11 and 12. These areas could be protected areas (PA) or Other Effective Area-Based Conservation Measures (OECMs), including indigenous reserves, private reserves, and any areas where extractive or productive activities are prohibited or regulated by voluntary schemes, certifications or law, in the interest of biodiversity conservation.

To identify potential regions in which to expand conservation areas to meet conservation objectives, we used the Marxan conservation planning software (Watts et al. 2009a). Marxan operates to design a near-optimal protected network of conservation areas which meets biodiversity targets while minimizing costs, *e.g.* opportunity costs, or management costs (Watts et al. 2009b). Our study region consisted of the global terrestrial extent, excluding Antarctica, divided into 2,063,413 grid cells each with a resolution of 5 arcmin (∼100 km^2^ at the Equator). Data on the current PA network (as of April 2019) was obtained from the World Database on Protected Areas (WDPA) downloadable from www.protectedplanet.net. Following Butchart et al. (2015), we excluded proposed protected areas, those with an unknown designation status, UNESCO biosphere reserves, and those lacking both reported extent and spatial boundaries. Cells with more than 50% of their area within the current PA network (281,701 cells, 13.7% of the study region) were considered as protected and locked into the planning solution. We adopted as a cost value, incurred when a cell is to be conserved in the solution generated by Marxan, the projected suitability values of each cell to agriculture in 2030, modelled by the Integrated Model to Assess the Global Environment (IMAGE) version 3.0 (Stehfest et al. 2014). IMAGE determines suitability following an empirical allocation algorithm with three drivers (Doelman et al. 2018): potential crop yield as modelled by LPJmL, accessibility, population density from the HYDE database (Goldewijk et al. 2010), and terrain slope index from the Harmonized World Soil Database (Nachtergaele et al. 2010).

We tested 3 Marxan prioritization scenarios using the three different targets: the RS, RSI and HSI scenarios. We considered targets as met when the conservation areas accounted for 99% of the target area for each species. For each scenario, Marxan was parameterized to perform 100 runs with 200,000,000 iterations in each. We used the best solution (the solution that meets most targets with minimal costs) of each scenario to (1) calculate the percentage of land to be conserved for the world and per continent, (2) calculate the contribution of the current PA network towards the RS, RSI and HSI targets (3) compare the IUCN-informed target setting method with that previously set by Rodrigues et al. (2004). For the latter, we calculated how much RSI and HSI targets were represented under the RS scenario conserved area network. Finally, we used cell selection frequency to (4) identify regions of highest conservation priority among the different scenarios. Therefore, we mapped all cells selected to be conserved in more than 90% of the runs, crucial towards fulfilling conservation objectives (Levin & Mazor 2015). We compared these regions among scenarios by overlaying high priority areas for range-based scenarios (RS and RSI) and for RL-informed targets (RSI and HSI). Finally, as areas with a higher agricultural potential are likely to be converted faster than unsuitable land, we overlaid the agricultural suitability values used as a cost layer in high priority areas for the RSI and HSI scenarios.

## RESULTS

### 1. Conservation targets

IUCN-informed area targets (RSI and HSI targets) are larger than RS targets for most species (93% of RSI targets > RS targets and 70% of HSI targets > RS targets). Targets based on suitable habitat (HSI targets) are either smaller than (80% of the targets) or equal to (20% of the targets, most of them amounting 2,200 km^2^ or 1,000,000 km^2^) RSI targets. We find 3% and 12% of species require the minimum protection of 2,200 km^2^ for the RSI and HSI targets respectively.

### 2. Conservation area networks

We find that 47% (2054 species), 9% (971 species), and 6% (248 species) of the species have their RS, RSI and HSI targets met within the current PA network, respectively. If we were only to consider threatened species (n=1197), these figures would be respectively 14.8% (178 species), 1.2% (15 species) and 0.7% (9 species). If we were to consider all threatened terrestrial vertebrates for which distributional ranges and habitat preferences were available (n=4720 among amphibians, birds and mammals), 9% (426 species) would be adequately protected when using RS targets, 1.06% (50 species) when using RSI targets and 0.5% (22 species) when considering HIS targets. An additional 4% of land suffices to meet all RS species targets for mammals (18% of the world’s terrestrial land, Fig. 1a) but would only allow to meet representation for 15% (670 species, RSI targets) and 8% (360 species, HSI targets) of the species’ IUCN informed targets. To meet most of these targets, 60% (RSI targets) to 62% (HSI targets) of the world’s terrestrial extent must be protected (Fig.1b and 1c). Substantial increase in conservation area coverage occur in Asia (almost 6 times the current coverage required), followed by Africa and North America (5 and 4 times the current coverage required). Despite having the highest current PA coverage, Oceania and South America require the highest proportion (>70%) of their land to be conserved to meet targets. RSI targets cannot be met for 66 species, for which known ranges fall short of the minimum target of 2,200 km^2^, and 435 species cannot meet their HSI targets due to the lack of available suitable land.

**Figure 1:**
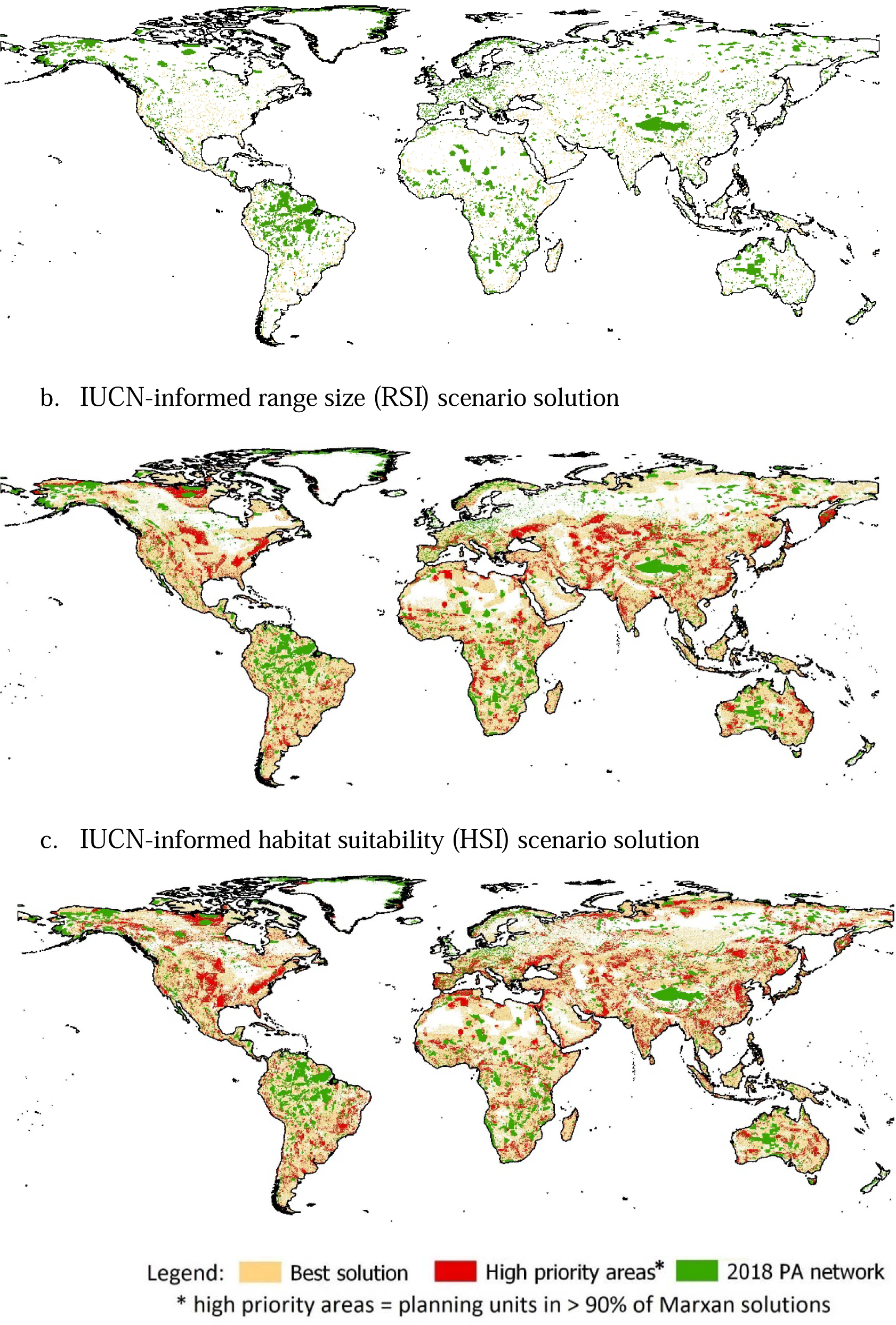
Conservation area networks generated as the best solution by Marxan with mammal species targets calculated based on A. range-size only (RS targets), B. IUCN-informed range-size only (RSI targets), and C. IUCN-informed habitat suitability only (HSI targets).

**Figure 2:**
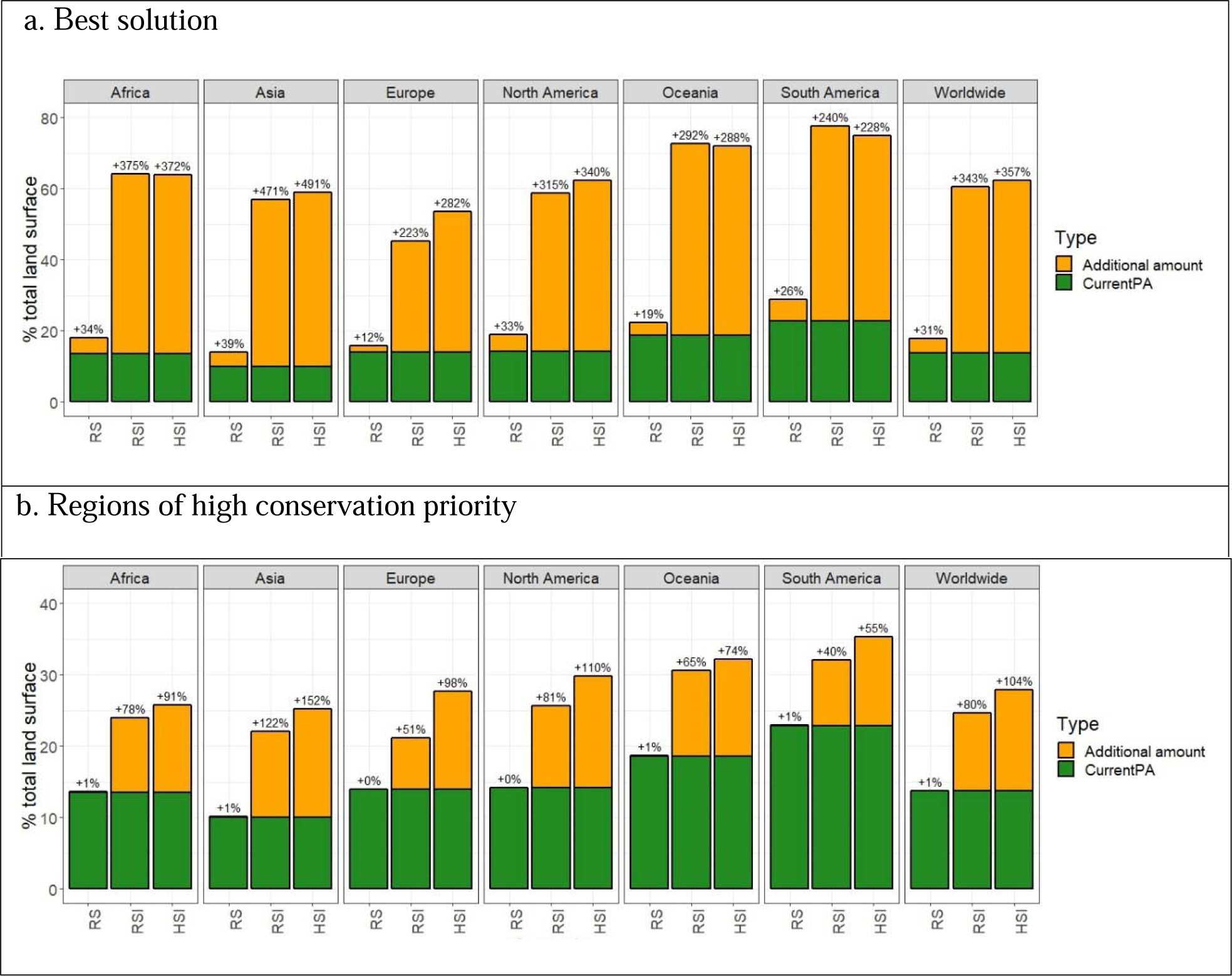
The required percentage increase of protected areas (WDPA, 2018) in each continent to meet: A. Marxan’s best solution, and B. all areas with a selection frequency greater than 90% across all Marxan runs for each scenario. Where Marxan scenarios are based on: RS, range-size targets (RS) only; RSI, IUCN-informed range-size targets only; HSI, IUCN-informed habitat suitability targets only.

High conservation priority areas cover 0.06% (RS), 11% (RSI) and 14% (HSI scenario) of the non-protected Earth’s surface, many of which are highly suitable for agriculture (Fig. S1a and S1b). Five percent of high priority areas overlap between RSI and HSI scenarios (Fig. 3). These areas are mainly located in North America (Appalachian range, mainland Nunavut, Dakota), in Asia (Middle East, Central Asia, Eastern Russian peninsulas, and Japan’s Ryukyu Islands), in Europe (Ukraine, around the Alps, Northern Spain and Southern Norway), in Africa (around the Tropics, in the Saharan Atlas and South Africa) and in Oceania (9% of Australia).

**Figure 3:**
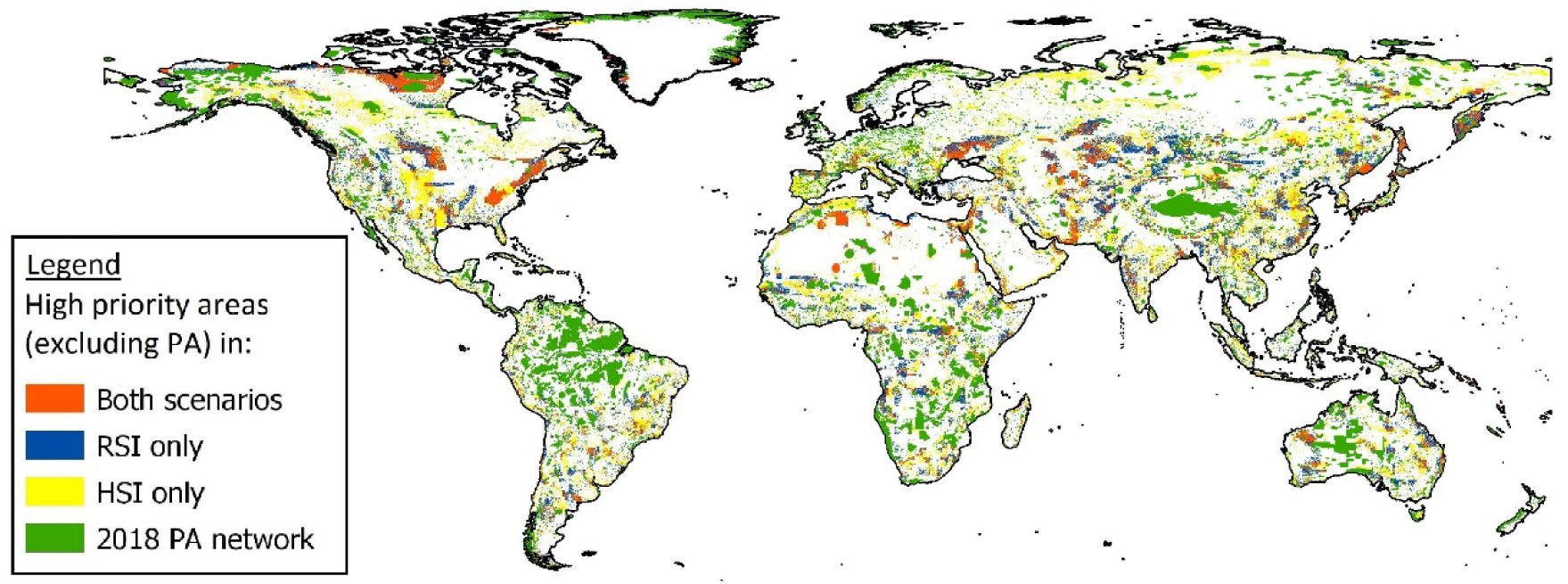
Regions of high conservation priority across RSI target scenarios and IUCN-informed habitat suitability only (HSI target) scenarios. Regions of high priority are those with a selection frequency of 90% or more across all Marxan runs for a scenario.

## DISCUSSION

Arbitrary range-based targets used in previous studies require 18% of the planet to achieve targets for all terrestrial mammal species considered here. However, implementing this network would leave more than 80% of terrestrial mammals at high risk of extinction. To ensure their persistence, at least 60% of the Earth’s surface (excluding Antarctica) must be managed to conserve biodiversity. Our results suggest that scientifically-based, bolder conservation targets will be needed to protect biodiversity in the future.

Despite an increase in coverage of the PA network to over 14% of the Earth’s land surface in 2018 (UNEP-WCMC & IUCN 2018), less than half of the range-based targets (RS targets) of mammal species are currently met. This confirms earlier findings that the rate of progress towards the protection of terrestrial mammal species has been disproportionally slower than the rate of increase in protected areas (Rodrigues et al. 2004; Butchart et al. 2015; Venter et al. 2018). Indeed the recent expansion of the PA network has privileged areas with low opportunity cost for agriculture and relatively low biodiversity value (Venter et al. 2018), thereby reducing the potential for protected areas to safeguard imperiled biodiversity.

To ensure the persistence of mammals, efforts to expand the PA network must be considerably more ambitious than the 17% prescribed by the Aichi Target 11 (CBD 2010). Our finding that 60% of the Earth’s surface must be managed to sustain biodiversity supports the idea that bold conservation targets and actions, such as those prescribed by the Half-Earth proposal, are urgently needed to guarantee a future for the planet’s biodiversity. The areas of high conservation priority identified in our scenarios provide specific guidance for future expansion of area-based conservation measures. Their protection, crucial to meet our conservation objectives, would require the expansion of the current PA network to twice its current size into areas that are sometimes highly suitable for agriculture and therefore likely to be rapidly converted in the future.

The use of habitat-based targets, compared to range-based targets, results in larger networks needed to protect fewer species. The use of ranges to evaluate species’ needs for conservation may thus result in optimistic estimates and may include areas where species are absent, and habitat is unsuitable for their reintroduction. Using suitable habitat to generate conservation targets constitutes a more ecologically meaningful representation of the actual distribution of the species and is more effective to design efficient protected area networks. However, using suitable habitat to set targets and inform conservation planning has its own limitation. Assuming ESH = AOO may be often invalid. Nonetheless, all priority areas where ESH ≠ AOO could be considered as candidate sites for reintroductions if after on-the-ground surveys, alternative sites where the species is still extant were not found and the conditions for reintroductions were favorable. In alternative, the value of these sites for the conservation of the species absent from the site, should be discounted and priorities reassessed.

To provide a more comprehensive account of the status of biodiversity and the progress achieved towards conservation objectives, more analyses of this type are needed. The first obvious step would be the inclusion of other taxa, especially those whose centers of endemism and high richness least overlap with mammals, e.g. plants or amphibians (Kier et al. 2009) to provide greater insight into the extent and spatial distribution of areas needing conservation efforts to minimize species extinction risk. Secondly, as prioritization is scale dependent, more localized analyses will be necessary, wherein connectivity between protected lands could be explicitly considered while new protected areas and OECMs could be included as they are created.

The urgent need to rapidly expand the current network of conserved areas to avoid extinctions and reduce the overall biodiversity decline, requires a collaborative, multidisciplinary international approach to avoid creation of “paper parks” without effective funding and management (Watson et al. 2014). This requires the strong involvement of stakeholders and empowerment of indigenous and local communities. Recognizing and integrating OECMs managed by these stakeholders may provide vital connective corridors between PAs, crucial to achieving adequate global biodiversity representation (CBD 2010; Locke 2013). Efforts should also be placed on protecting disconnected populations and favoring recolonization of lost habitat to reconnect them. Many of the species for which our analyses failed to meet the targets are island species, mainly located around South-East Asia, or the Japanese Ryukyu archipelago. Habitat restoration or species re-introduction may be viable conservation options if a species has recently become locally extinct from islands once within its historical range.

While necessary to achieve the protection of terrestrial mammal species, conserving over 60% of the terrestrial surface is a highly ambitious target, requiring an extensive multidisciplinary, internationally coordinated approach, which may take years to fully implement. In the short-term, initial conservation efforts should be focused on protecting high priority areas, such as those highlighted in the HSI scenario, and that are facing the most imminent threat of conversion or degradation. A carefully curated and coordinated expansion of the current PA network to 17% coverage could potentially triple the average species and ecoregion coverage (Pouzols et al. 2014). Going forwards, cooperation across international scales, as well as involving key local stakeholders such as indigenous peoples will be vital to effectively implement a global and comprehensive network of interconnected PAs and OECMs.

## SUPPORTING INFORMATION

### Box 1: ICUN Red List categories and criteria

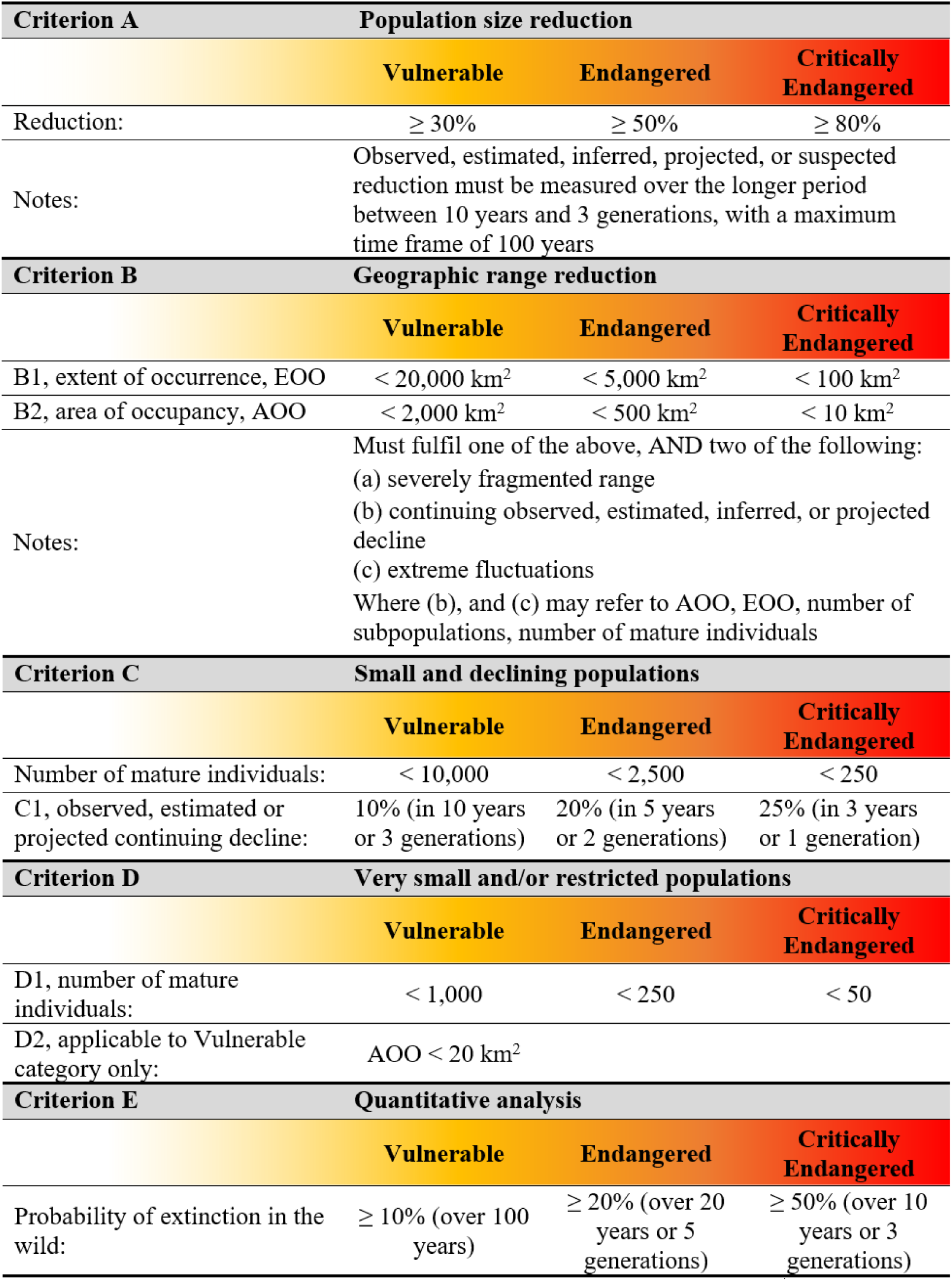

**FIGURE S1.**
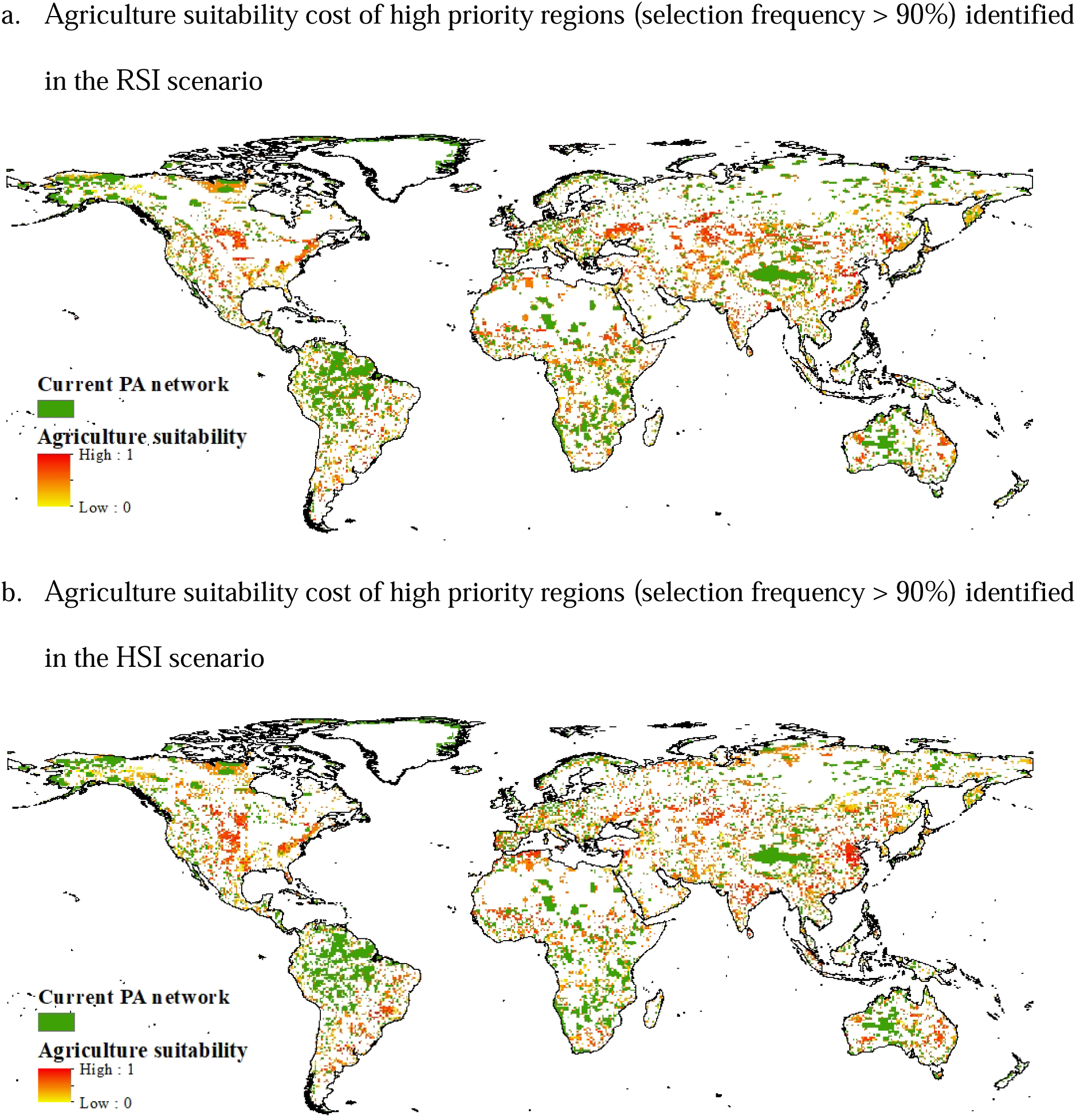

